# No evidence sex-specific senescence varies with resource competition in Soay sheep

**DOI:** 10.64898/2026.09.02.748798

**Authors:** Elizabeth D. Drake, Xavier Bal, Jill G. Pilkington, Josephine M. Pemberton, Daniel H. Nussey, Hannah Froy

**Affiliations:** Institute of Ecology and Evolution, School of Biological Sciences, University of Edinburgh, Edinburgh, EH9 3FL, UK

**Keywords:** Ageing, density dependence, environmental sensitivity, physiological resilience, selective disappearance, life-history evolution

## Abstract

Demographic senescence, the age-related decline in survival and reproduction, is widespread. Older individuals, and particularly males in polygynous systems, are expected to be disproportionately vulnerable to environmental stressors due to physiological decline and high reproductive costs. However, studies testing if harsh conditions exacerbate senescence, and whether this effect is sex-dependent, remain limited in wild populations. Using 40 years of data from wild Soay sheep (*Ovis aries*) on St Kilda, Scotland, we tested whether increased resource competition (indexed by population density) accelerates demographic senescence, and if late-life environmental sensitivity is male-biased. High densities consistently suppressed overall adult performance, making individuals less likely to survive, breed, or rear high- quality offspring under more intense competition. Although survival and reproductive traits exhibited clear senescence, increased population density did not accelerate late- life declines in any trait. Rates of decline with age were unaffected by density and did not differ between the sexes. Our results challenge the assumption that older individuals and males are inherently more vulnerable to harsh conditions, suggesting physiological or behavioural buffering may maintain late-life resilience against resource limitation in some systems.

## Introduction

Demographic senescence, the age-related decline in survival and reproductive performance, is widespread and plays an important role in ecological and evolutionary dynamics (Nussey *et al*., 2013; Jones *et al*., 2014; Mitchell *et al*., 2015). While senescence rates vary considerably among species and populations, how environmental conditions and sex independently and interactively shape demographic senescence in wild animals remains poorly understood (Jones *et al*., 2014; Gaillard & Lemaître, 2020). Increasing empirical evidence suggests deteriorating physiological function with age increases vulnerability to environmental stress, such that older individuals may experience more rapid demographic decline under harsh current conditions (Williams *et al*., 2006; Reichert *et al*., 2010; Pardo *et al*., 2013; Holand *et al*., 2016). Evolutionary theory further predicts sex differences in lifespan, senescence and environmental sensitivity in polygynous species with strong male-male competition and marked sexual dimorphism (Maklakov & Lummaa, 2013; Brooks & Garratt, 2017). Although males in such systems are generally shorter lived and more demographically sensitive to early life environmental conditions than females (e.g., Kruuk *et al*., 1999; Hamel *et al*., 2016; Staerk *et al*., 2025), evidence for faster male senescence remains scarce. Crucially, the prediction that demographic senescence is more sensitive to environmental pressures in males than females has yet to be tested in a natural population.

Bio-gerontological research increasingly supports the idea that declining biological function in later life reduces an organism’s physiological resilience: its ability to withstand or recover from environmental challenges (Promislow *et al*., 2022). Under this hypothesis, demographic senescence is at least partly driven by the failure of physiological resilience and should be exacerbated under more challenging environmental conditions. Senescence of physiological traits such as body condition, immune response or foraging ability may result in decreased physiological resilience and thus increased demographic sensitivity to adverse current conditions. There is emerging evidence that senescence is exacerbated under challenging environmental conditions in elderly humans (Lee *et al*., 2022), experimental laboratory organisms (Dudycha, 2003; Briga *et al*., 2017; Sanghvi *et al*., 2022) and wild vertebrates (Mysterud *et al*., 2001; Reichert *et al*., 2010; Pardo *et al*., 2013).

Comparative studies strongly support the expectation from evolutionary theory that, in polygynous mammals, females are longer lived than males (Tidière *et al*., 2015; Lemaître *et al*., 2020; Staerk *et al*., 2025). How and why males and females differ in the onset and rate of demographic senescence remains less well resolved in wild mammal populations (Maklakov & Lummaa, 2013; Lemaître *et al*., 2020; Bronikowski *et al*., 2022). Classical evolutionary theory predicts males should evolve faster senescence than females when their risk of adult mortality from environmental causes is higher (Williams, 1957). More recent theory also suggests sex-biased ageing rates may evolve as a consequence of the accumulation of mutations in genes with sex-biased fitness effects (Moorad *et al*., preprint). While empirical studies have reported sex differences in age-specific survival, reproductive traits and body mass in several wild populations (e.g., Murgatroyd *et al*., 2018; Macdonald *et al*., 2025; reviewed in Bronikowski *et al*., 2022), a large-scale comparative study found no consistent actuarial senescence bias between the sexes across mammals (Lemaître *et al*., 2020). This work highlighted that sex-specific senescence may be difficult to generalise across heterogeneous ecological conditions experienced by wild populations.

In polygynous species, the costs of male secondary sexual traits and damaging intrasexual competition leads to the expectation that males should be more sensitive to environmental pressures than females (Maklakov & Lummaa, 2013; Brooks & Garratt, 2017). Adult lifespan is more sensitive to early life adversity in males than females in wild mammals (e.g., Solberg *et al*., 2004; Tidière *et al*., 2015; Drake *et al*., 2025). However, much less attention has been paid to differences in the sensitivity to environmental pressures during later life as a driver of sex differences in demographic senescence. Recent studies have shown sex differences in the environmental sensitivity in actuarial senescence in wild anurans (Cayuela *et al*., 2021) and in body mass senescence in wild red deer (Mysterud *et al*., 2001). Lemaitre *et al*. (2020) suggested consistent male-biased actuarial senescence among mammals might only appear under challenging environmental conditions, noting male-biased senescence in bighorn sheep populations facing harsh winters but not in those with consistent resource availability. Their hypothesis, that in polygynous species, demographic senescence should be more environmentally sensitive in males than females, remains to be directly tested in a wild mammal.

Here, we use data from a long-term study of Soay sheep (*Ovis aries*) on the St Kilda archipelago, Scotland to test two hypotheses: (1) demographic senescence is faster under challenging environmental conditions, and (2) this late-life environmental sensitivity is male-biased. Actuarial senescence from around 6 years onwards has been documented in this population in both sexes (Catchpole *et al*., 2000; Colchero & Clark, 2012). Females experience reproductive senescence in annual breeding probability, offspring birth weight, and offspring recruitment success, but not twinning probability (Hayward *et al*., 2013, 2015). Previous studies suggest male weight, testes size and testosterone levels decline from around six years of age, but found no statistical support for a later-life decline in annual reproductive success (Preston *et al*., 2012; Hayward *et al*., 2015). We focus on annual population size as our focal measure of environmental conditions, as there is good evidence high population size reflects increased competition for food and mates, with well-established negative consequences for survival and reproductive performance in this system (Catchpole *et al*., 2000; Coulson *et al*., 2001; Jones *et al*., 2005).

## Methods

### Study system

Soay sheep (*Ovis aries*) have lived unmanaged on the St Kilda archipelago, off the West coast of Scotland, for millennia. The Village Bay population on the largest island, Hirta, has been individually monitored since 1985. Lambs are born in the spring (April-May); over 90% of lambs born are caught soon after birth, weighed and ear tagged for individual identification (Clutton-Brock & Pemberton, 2004). Most are singletons, although ∼18% of births are twins. The annual rut occurs between late October and early December, when males compete for access to oestrus females. Mortality peaks in late winter/early spring (February-April) and is male- and juvenile-biased. During three annual field expeditions, ten population censuses are performed within the study area. Mortality searches are conducted during the peak mortality period and recover ∼80% of deceased sheep. All data collection was conducted under UK Home Office license (current license: PP4825594).

The sheep in Village Bay experience pronounced inter-annual population size fluctuations (211-672 individuals), driven by density dependence and winter weather (Clutton-Brock *et al*., 1992; Coulson *et al*., 2001). High population size increases resource competition and parasite burdens, which depresses over-winter survival, and limits both male and female reproductive success (Clutton-Brock *et al*., 1991; Milner *et al*., 1999; Catchpole *et al*., 2000; Coulson *et al*., 2001).

### Data analysis

We modelled age-related changes in survival and reproductive success using data collected between 1985 and 2024. Demographic performance typically increases over early life, plateaus and then declines (demographic senescence) in wild vertebrates. We sought to exclude data from early ages when demographic performance was still increasing. Visual inspection of age-specific survival and reproductive performance traits suggested this was best achieved by including ages 3 and older for survival and 4 and older for reproductive traits (Figure S1). To model age-related changes in survival and reproductive performance, we used threshold (piecewise) regression models. These allow regression slopes to vary independently either side of a break point. Although quadratic functions are often used to model demographic ageing patterns in wild animals, these assume symmetrical changes either side of a peak. This assumption has no clear biological basis, and threshold functions allow us to test for independent demographic plateau and declines phases during adulthood (Toms & Lesperance, 2003; Descamps *et al*., 2008; Froy *et al*., 2013).

### Measures of survival, reproductive performance and population size

Annual survival (from May year *t* to May year *t+1)* was modelled as a binary trait (1=survived, 0=died; Figure 1). We analysed 4701 annual survival measurements from 1020 individuals. Death year and month was known for the vast majority of these individuals. For a small subset of individuals (146 of 1020 individuals), death year was known but month was not known precisely. Here, it was estimated using census data to identify when an individual was last sighted, assuming the animal survived if it was seen alive in a census after 1 May in the year of recorded death (21 individuals). If it was not seen after this point it was assumed to have died over that winter period (125 individuals).

**Figure 1.**
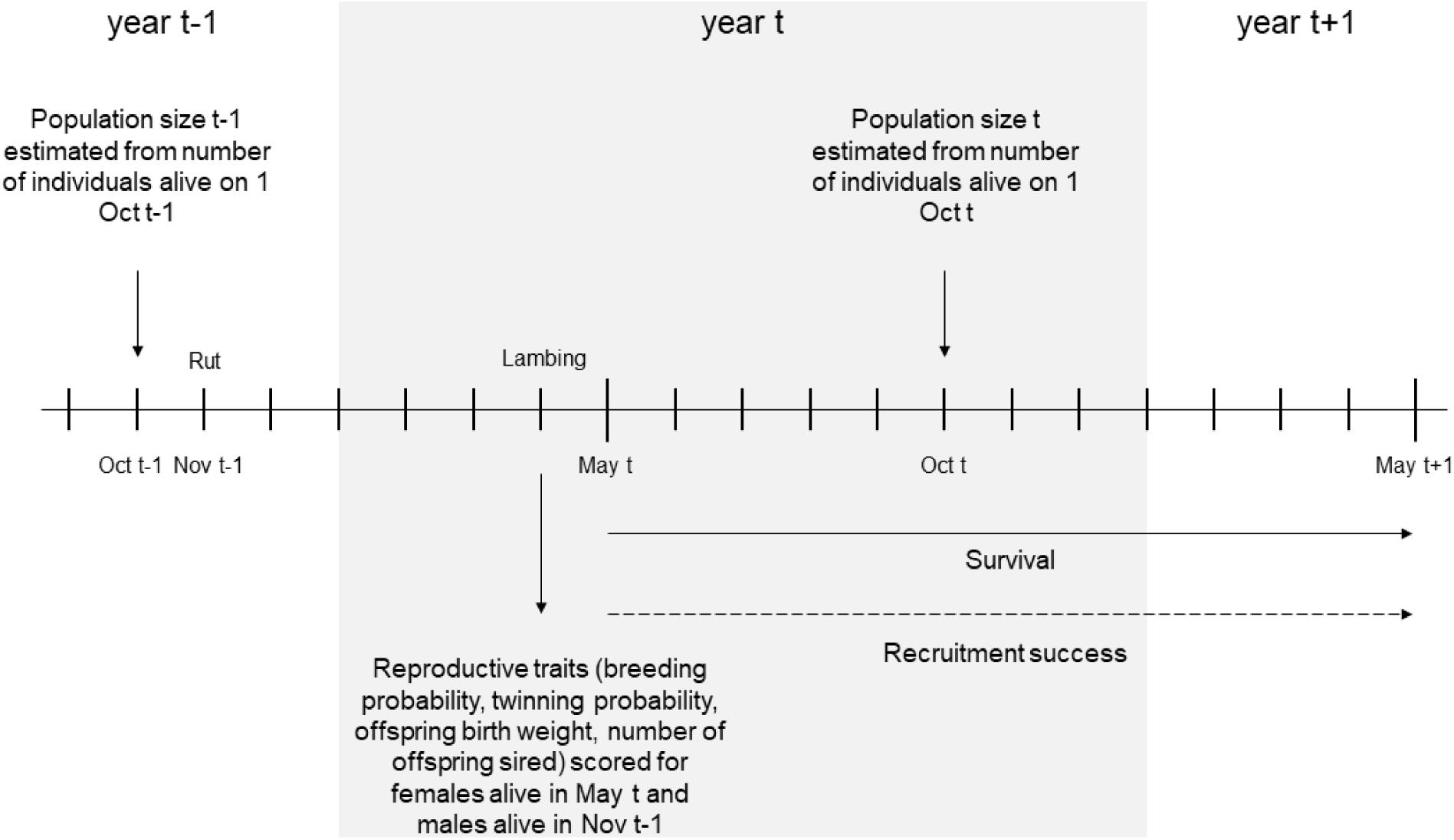
Timeline of when population size measures are estimated and, survival and reproductive traits are scored. In analyses, population size in year t-1 is related to reproductive traits (breeding probability, twinning probability, offspring birth weight, number of offspring sired) in year t, and population size in year t is related to survival and recruitment success from May t to May t+1.

To capture the multiple aspects of annual reproductive success, we decomposed this trait into annual breeding probability and four sex-specific breeding components that were conditional on having bred (female traits: twinning probability, offspring birth weight and offspring recruitment success; male traits: number of offspring sired). Survival and breeding probability were the only traits where male and female slopes could be directly compared.

Breeding probability was a binary trait (1=individual did give birth to or sire an offspring in year t, 0= individual did not), assigned for males present in the Village Bay area during the preceding rut and females alive on 1 May year t (i.e., females that survived lambing), to avoid conflating winter mortality and breeding success (N=5061 observations of 1225 individuals; Figure 1). Breeding probability assignment differed between the sexes as males do not have to be alive in the spring in order to sire an offspring, but females do. Twinning probability was a binary trait indicating whether a female gave birth to twins in year t, providing she produced at least one lamb (Figure 1; N=3155 observations of 737 females). Recruitment success was a binary trait indicating whether the offspring of a given female born in year t survived until 1 May year t+1 (Figure 1; N=3638 observations of 738 females). Offspring birth weight was a continuous trait in kilograms for offspring born in year t and weighed within the first seven days of life (N=2881 observations of 707 females). The age of offspring in days was included as a fixed effect in models of offspring birth weight to account for rapid growth during this early period. Birth weight data from the 2001 and 2020 cohorts are not included in our analyses as logistical challenges resulted in a high proportion of missing data (foot and mouth outbreak and Covid-19 pandemic, respectively). Number of offspring sired was the number of live offspring born in year t (ranging 1–22), conditional on the male having sired at least one lamb that year (Figure 1; N=648 observations of 256 males). For both sexes, reproductive performance was calculated based on a multigenerational genetic pedigree, inferred using a subset of 431 unlinked Single Nucleotide Polymorphisms (SNPs) derived from the Illumina Ovine 50K SNP array using the R package Sequoia (Bérénos *et al*., 2014; Huisman, 2017). For a small number of cases where SNP genotype information was not available, parentage assignments were made using field observations (for females: 11.2%) or from microsatellite data (for males: 3.0%; Morrissey *et al*., 2012).

Population density, our index of resource competition, was estimated as the number of individuals alive on 1 October (mean=480.722, SD=112.585). Survival and recruitment success were related to population size in year t, and all other reproductive traits were related to population size in year t-1 (Figure 1).

### Data analysis

Annual survival, breeding probability, twinning probability and recruitment success were modelled as binomial generalised linear mixed models (GLMMs; logit link); male annual number of offspring sired as a negative binomial GLMM (log link); and offspring birth weight as a Gaussian linear mixed model. All models included individual identity and measurement year as random intercept terms to account for repeated measurements of individuals and environmental conditions. A cohort-level random effect was retained in only the survival models, as the variance estimate for cohort was very small (<0.001 in the majority of cases) and the confidence intervals very wide (0.000-Inf in all cases) in every reproductive trait model.

To determine the best age function for each trait, we compared a linear age term or a single age threshold that varied between 3 and 8 years for survival and 4 and 9 years for the reproductive traits (higher thresholds were not tested as there were <30 observations per age per sex beyond 8 and 9 years, respectively). Separate models of each sex were also run, testing the same age functions, to determine if using the same threshold for both sexes was appropriate (Table S2). Survival and breeding probability models included a fixed effect of sex to account for differences in average trait values between the sexes, and an age-by-sex interaction to allow the ageing slopes to differ between males and females. The best age function model was selected based on an AIC difference of ≤2 compared to the lowest-AIC model. Threshold confidence intervals were estimated by varying breakpoints by 0.01 increments and identifying where the ΔAIC score was ≥2 relative to the best age function model.

To account for selective disappearance, age at last observation was included in reproductive models (van de Pol *et al*., 2006). A last-recorded-age-by-sex interaction was included in the breeding probability model to allow for differing selective disappearance between the sexes. Offspring sex was included as a fixed effect in the models of offspring birth weight and offspring recruitment, as male lambs are known to be larger and have reduced first-year survival probability (Clutton-Brock *et al*., 1992; Drake *et al*., 2025).

After determining the best age function for each trait, we moved to testing the effects of population size on age-related changes in survival and reproductive performance. First, population size in year t-1 was added as a fixed covariate to the best age function models of breeding probability, twinning probability, offspring birth weight, and number of offspring sired, and population size in year t was added as a fixed covariate to the best age function models of survival and recruitment probability (Figure 1). Then, an interaction between age and the relevant measure of population size was added to all models. Interactions between sex and age, and sex and the relevant measure of population size were also added to the models of survival and breeding probability, to aid with the interpretation of the final models. Finally, to test for sex-specific environmental effects on ageing, a three-way between age, sex, and the relevant measure of population size was added to the models of survival and breeding probability. Population size was scaled to mean = 0, standard deviation = 1 prior to inclusion in the models to facilitate interpretation of interaction terms. The AICs of all models run per trait were compared to aid interpretation of results.

All analyses were performed using R (R Core Team, 2026). Models were run using the package *glmmTMB 1.1.11* (Brooks *et al*., 2017). Models of offspring birth weight comparing different age functions were run using Maximum Likelihood Estimation but the final models were estimated using Restricted Maximum Likelihood. Variance and 95% confidence intervals for random effects were calculated using the package *stats* (R Core Team, 2026). Model fit was validated using *DHARMa 0.4.7* (Hartig *et al*., 2024), and plots were generated using *ggplot2 4.0.0* (Wickham, 2016) and *ggeffects 2.3.1* (Lüdecke, 2018).

## Results

### Survival senescence

The best age function model for annual survival probability did not include a threshold age term (Table S1A). Survival declined after age 3 in both sexes (female slope: - 0.440±0.042, p<0.001; male slope: -0.402±0.069, p<0.001; Table S3A; Figure 2A).

**Figure 2.**
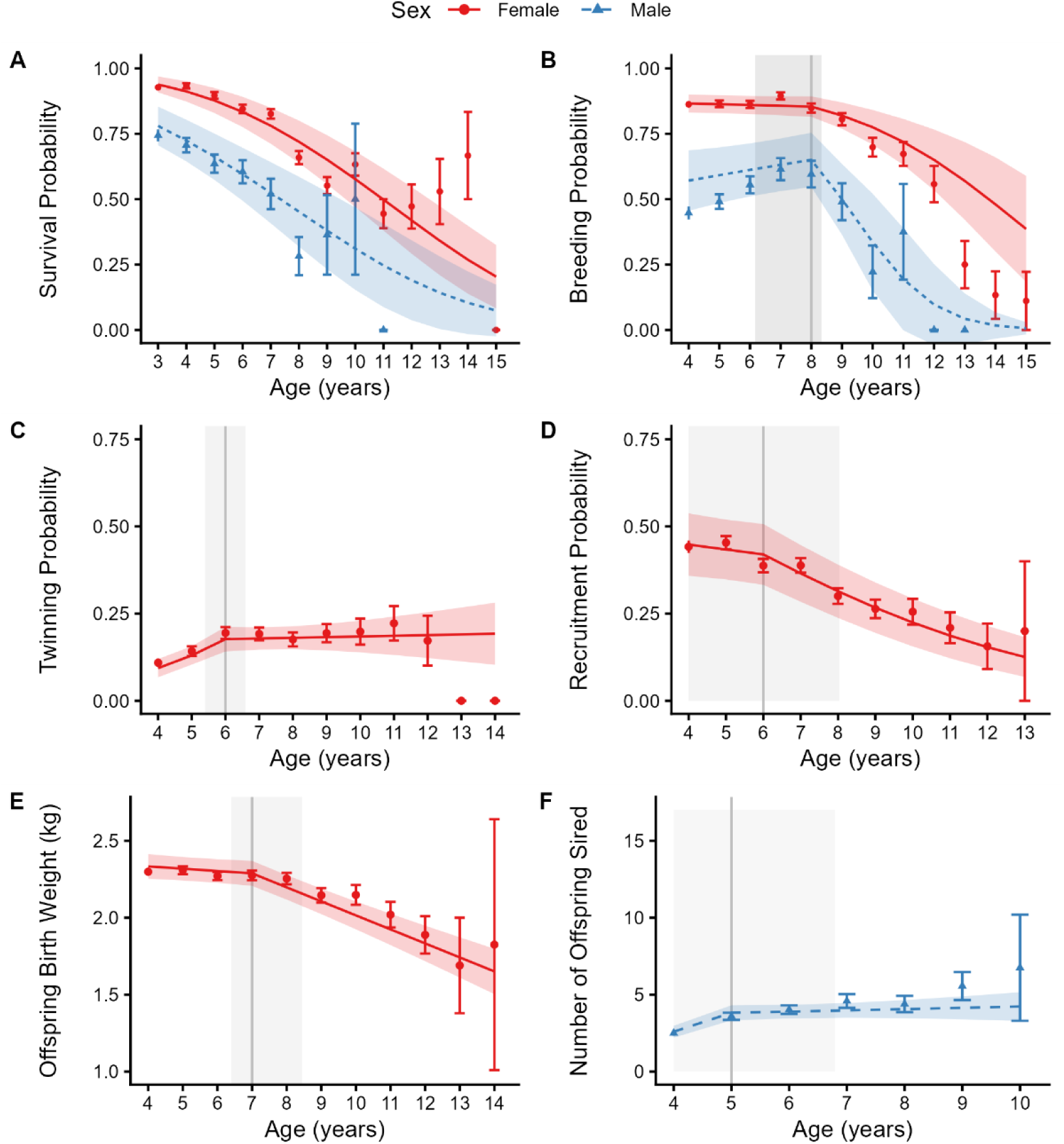
Estimates of the effect of age from the best age function models for both females and males from models of (A) survival probability and (B) breeding probability; for female only models of (C) twinning probability, (D) recruitment probability, (E) offspring birth weight; and (F) from a model of number of offspring sired by males in Soay sheep. Red circular and blue triangular points and error bars represent the mean and associated standard error at each age for females and males, respectively. Red solid and blue dashed lines and ribbons represent predictions and the 95% confidence intervals for the model estimates for females and males respectively. Model estimates are from the best age function model for each trait and can be found in tables S3A, S4A, S5 respectively. Grey solid vertical lines and shading represent thresholds and associated 95% confidence intervals.

Females survived consistently better than males, but senescence rates did not differ between the sexes (Age:Sex interaction: 0.038±0.057, p=0.505; Table S3A; Figure 2A). Adding a fixed covariate of population size showed that high population size significantly reduced overall survival (-0.641±0.233, p=0.006; Tables 2A&S3B). All two-way and three-way interactions among age, sex, and density were non-significant, suggesting the effect of population size did not vary between the sexes or depending on age and that sex-differences in survival senescence are not exacerbated under high population densities (Tables 1 & S3C; Figure 3A). Adding the interactions between age, sex, and population size did not improve model fit (Table 2A).

**Figure 3.**
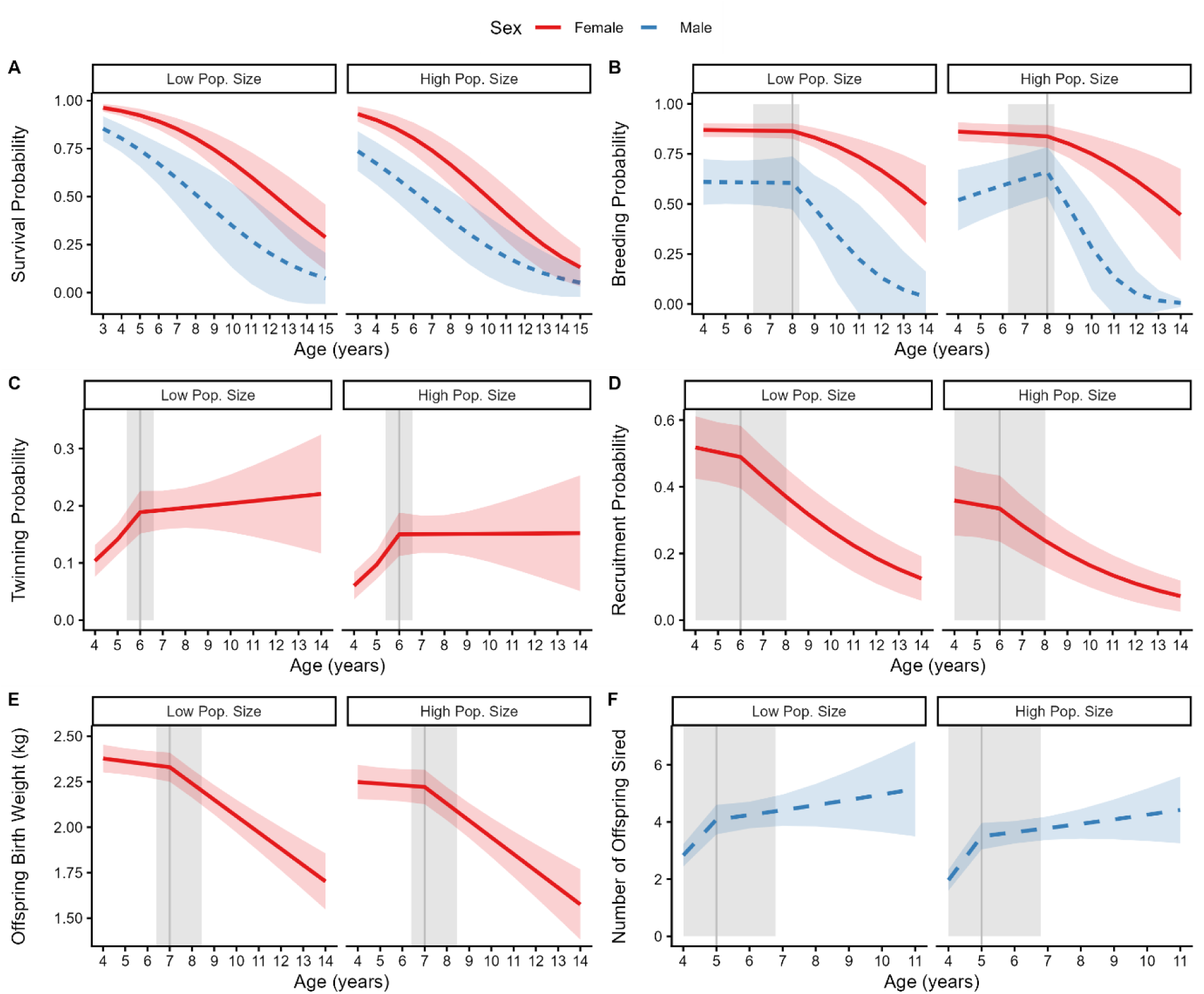
Estimates of the effect of age and population size on male and female (A) survival probability and (B) breeding probability; female (C) twinning probability, (D) recruitment probability, (E) offspring birth weight; and (F) the number of offspring sired by males in Soay sheep. Red and blue lines and ribbons represent predictions and the 95% confidence intervals for the model estimates for females and males respectively (Table 1, 2&3). Grey lines and ribbons represent thresholds and the associated 95% confidence intervals. For the low and high population size plots, the population size covariate was held at the 25% (447 individuals) and 75% (580 individuals) quantile for population size respectively

**Table 1.** . Summary of the fixed and random effects of a binomial GLMM of survival, with a three-way interaction between age, sex and population size in year t. Reference level for Sex is female. Significant fixed effects in bold. Nfemale=3604 observations of 649 individuals; Nmale=1097 observations of 371 individuals.

| Fixed effects | Estimate | Std. error | p-value | Random effects | Variance | 95% CI |
| --- | --- | --- | --- | --- | --- | --- |
| **Intercept** | **4.446** | **0.362** | **<0.001** | Identity | 0.078 | 0.002-2.497 |
| **Sex (male)** | **-1.770** | **0.320** | **<0.001** | Birth year | 0.019 | 0.002-0.163 |
| **Age** | **-0.425** | **0.042** | **<0.001** | Measurement year | 1.754 | 0.992-3.100 |
| Population size | -0.472 | 0.285 | 0.098 |  |  |  |
| Age:Sex (male) | 0.022 | 0.061 | 0.717 |  |  |  |
| Age:Population size (year $t$) | -0.027 | 0.027 | 0.315 | | | |
| Sex (male):Population size (year $t$) | -0.318 | 0.331 | 0.337 | | | |
| Age:Sex (male):Population size (year $t$) | 0.059 | 0.064 | 0.358 | | | |

**Table 2.** AICs of models of (A) survival, (B) breeding probability, (C) twinning probability, (D) offspring birth weight, (E) recruitment probability, and (F) number of offspring sired, with fixed covariate of the relevant population size measure and various interactions between age, sex, and population size. Population size in year t (PSt), Population size in year t-1 (PSt-1), last recorded age (LRA)..

| Model | AIC | ΔAIC |
| --- | --- | --- |
| A) Survival |  |  |
| Best age function model (I + Age + Sex + Sex:Age) | 3517.784 | 4.896 |
| **Best age function model + $PS_t$** | **3512.888** | **0.000** |
| Best age function model + $PS_t$ + Age:$PS_t$ + Sex:$PS_t$ | 3516.321 | 3.433 |
| Best age function model + $PS_t$ + Age:$PS_t$ + Sex:$PS_t$ + Age:Sex:$PS_t$ | 3517.478 | 4.590 |
| B) Breeding probability |  |  |
| **Best age function model (I + Age + Sex + LRA + Age:Sex + Sex:LRA)** | **3518.617** | **0.000** |
| Best age function model + $PS_{t-1}$ | 3519.897 | 1.280 |
| Best age function model + $PS_{t-1}$ + Age:$PS_{t-1}$ + Sex:$PS_{t-1}$ | 3525.518 | 6.901 |
| Best age function model + $PS_{t-1}$ + Age:$PS_{t-1}$ + Sex:$PS_{t-1}$ + Age:Sex:$PS_{t-1}$ | 3523.251 | 4.634 |
| C) Twinning probability |  |  |
| Best age function model (I + Age + LRA) | 2570.089 | 17.536 |
| **Best age function model + $PS_{t-1}$** | **2552.553** | **0.000** |
| Best age function model + $PS_{t-1}$ + Age:$PS_{t-1}$ | 2554.423 | 1.870 |
| D) Recruitment probability |  |  |
| Best age function model (I + Age + LRA + Offspring sex) | 3841.655 | 6.387 |
| **Best age function model + $PS_t$** | **3835.268** | **0.000** |
| Best age function model + $PS_t$ + Age:$PS_t$ | 3839.262 | 3.994 |
| E) Offspring birth weight |  |  |
| Best age function model (I + Age + LRA + Offspring sex + Offspring capture age) | 4211.861 | 6.204 |
| **Best age function model + PS<sub>t-1</sub>** | **4205.657** | **0.000** |
| Best age function model + PS<sub>t-1</sub> + Age:PS<sub>t-1</sub> | 4209.322 | 2.539 |
| F) Number of offspring sired |  |  |
| Best age function model (I + Age + LRA) | 2808.686 | 20.385 |
| Best age function model + PS<sub>t-1</sub> | 2790.434 | 2.133 |
| **Best age function model + PS<sub>t-1</sub> + Age:PS<sub>t-1</sub>** | **2788.301** | **0.000** |

### Both sexes: Breeding probability senescence

The best age function model for breeding probability included a threshold at age 8 (CI 6.17-8.32; Table S1B), with single-sex best age function models confirming similar breakpoints (females: 7 (6.00-8.27); males: 8 (5.52-8.44); Table S2). Before age 8, age trajectories differed significantly between the sexes (Age_<8_:Sex interaction: 0.195±0.094, p=0.038; Table S4A). Female breeding probability remained stable with age (-0.056±0.057, p=0.327), whereas males breeding probability showed a marginally non-significant increase with age (0.140±0.076, p=0.067; Figure 2B; Table S4A). After age 8, breeding probability declined significantly in both sexes (female age slope: -0.537±0.078, p <0.001; male age slope: -1.022±0.287, p<0.001; Figure 2B; Table S4A). However, there was no evidence the rate of senescence differed between the sexes (Age_>8_:Sex interaction: -0.497±0.287, p=0.084; Table S4A). Males who reached older last recorded ages had higher average breeding probabilities compared to those who died younger, but there was no evidence of selective disappearance among females (Table S4A; male last recorded age slope: 0.365±0.082, p <0.001).

After adding a fixed covariate of population size and the interactions between age, sex, and population size, there was a significant two-way interaction between sex and population size. There was a significant effect of population size on only male breeding probability (-1.157±0.483, p=0.017; Table 3), suggesting males performed worse than females at higher population densities. The three-way interaction between age, sex, and population size was significant only prior to the threshold at age 8, suggesting younger males performed relatively worse than females at higher population densities (Table 3; Figure 3B). There was no statistical support for age-dependent effects of population size on breeding probability in later life in either sex, and no evidence sex differences in senescence varied depending on population density (Figure 3; Table 3). Adding the interactions between age, sex, and population size did not significantly improve the fit of this model (Table 2B).

**Table 3.**
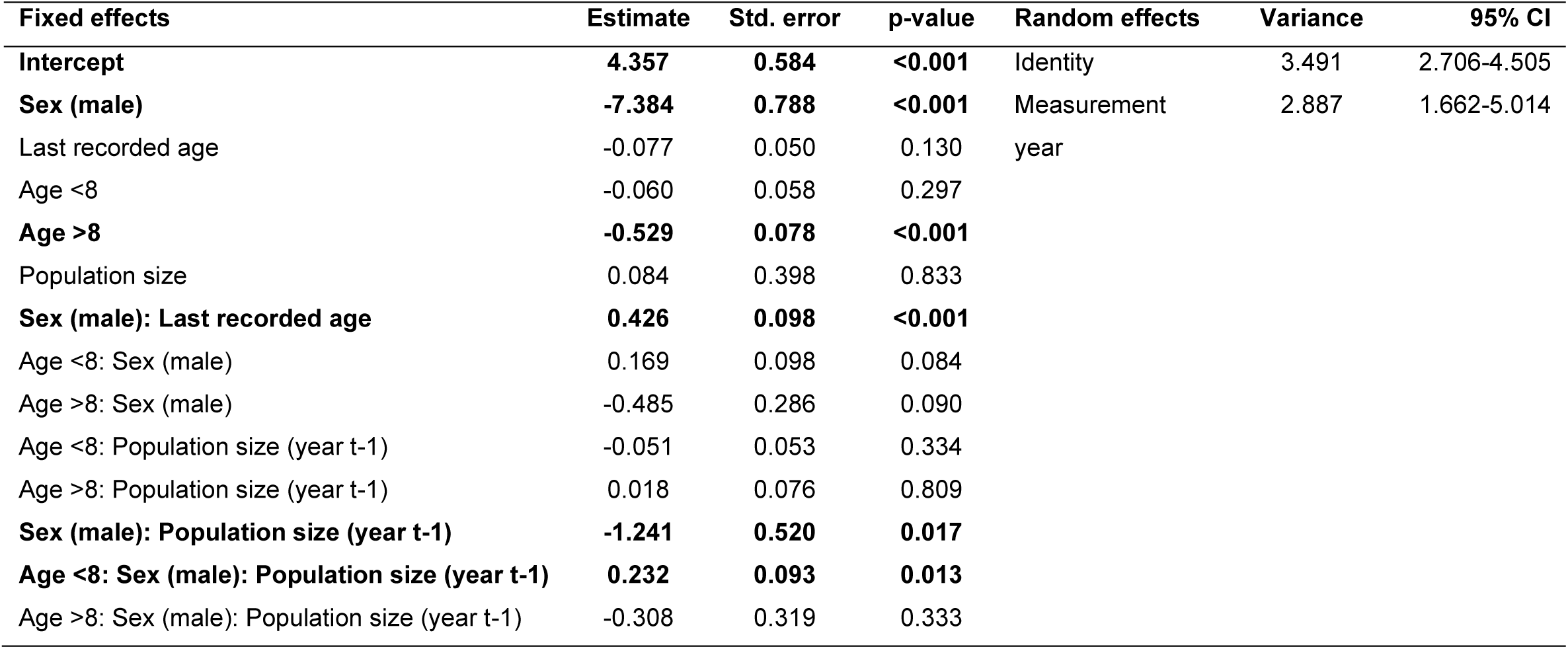
. Summary of the fixed and random effects of a binomial GLMM of breeding probability, with a threshold at age 8 with a three-way interaction between age, sex and population size in year t-1. Reference level for Sex is female. Significant fixed effects in bold. Nfemale=3774 observations of 807 individuals; Nmale=1287 observations of 418 individuals.

### Females: Twinning probability

The best age function model had a threshold at age 6 (CI 5.40-6.60; Table S1C). Twinning probability increased until age 6 and remained stable afterwards (Age_>6_ slope: 0.016±0.050, p=0.753; Table S5A; Figure 2C). Higher population densities decreased overall twinning probability (-0.339±0.077, p<0.001; Table S6A), however there was no significant interaction between age and population size (Table 4A; Figure 3C). The model with a fixed covariate of population size was the best fitting model for twinning probability (Table 2C).

**Table 4.** Summary of the fixed and random effects of GLMMs with an interaction between age and population size for (A) twinning probability (binomial; threshold at age 6), (B) recruitment success (binomial; threshold at age 6; reference level female offspring), (C) offspring birth weight (gaussian; threshold at age 7; reference level female offspring) and (D) number of offspring sired by a male (negative binomial; threshold age at 5). Significant fixed effects in bold.

| Fixed effects | Estimate | Std. error | p-value | Random effects | Variance | 95% CI |
| --- | --- | --- | --- | --- | --- | --- |
| A) Twinning probability, N=3155 observations of 737 females |  |  |  |  |  |  |
| **Intercept** | **-4.751** | **0.573** | **<0.001** | Identity | 2.246 | 1.660-3.041 |
| Last recorded age | -0.026 | 0.044 | 0.555 | Measurement | 0.062 | 0.020-0.188 |
| **Age <6** | **0.482** | **0.092** | **<0.001** | year |  |  |
| Age >6 | 0.020 | 0.050 | 0.696 |  |  |  |
| **Population size (year t-1)** | **-0.994** | **0.461** | **0.031** |  |  |  |
| Age <6: Population size (year t-1) | 0.125 | 0.088 | 0.155 |  |  |  |
| Age >6: Population size (year t-1) | -0.020 | 0.050 | 0.682 |  |  |  |
| B) Recruitment success, N=3638 observations of 738 females |  |  |  |  |  |  |
| Intercept | 0.060 | 0.465 | 0.898 | Identity | 0.625 | 0.435-0.897 |
| **Last recorded age** | **0.095** | **0.029** | **0.001** | Measurement | 1.468 | 0.876-2.461 |
| **Offspring sex (male)** | **-0.661** | **0.090** | **<0.001** | year |  |  |
| Age <6 | -0.071 | 0.065 | 0.270 |  |  |  |
| **Age >6** | **-0.313** | **0.043** | **<0.001** |  |  |  |
| Population size (year t) | -0.614 | 0.387 | 0.112 |  |  |  |
| Age <6: Population size (year t) | 0.003 | 0.067 | 0.963 |  |  |  |
| Age >6: Population size (year t) | 0.001 | 0.044 | 0.974 |  |  |  |
| C) Offspring birth weight, N=2881 observations of 707 females |  |  |  |  |  |  |
| **Intercept** | **1.763** | **0.086** | **<0.001** | Identity | 0.092 | 0.078-0.109 |
| **Last recorded age** | **0.016** | **0.007** | **0.026** | Measurement | 0.031 | 0.018-0.054 |
| **Offspring capture age (days)** | **0.115** | **0.004** | **<0.001** | year |  |  |
| **Offspring sex (male)** | **0.093** | **0.018** | **<0.001** |  |  |  |
| Age <7 | -0.013 | 0.009 | 0.138 |  |  |  |
| **Age >7** | **-0.091** | **0.010** | **<0.001** |  |  |  |
| **Population size (year t-1)** | **-0.117** | **0.056** | **0.036** |  |  |  |
| Age <7: Population size (year t-1) | 0.005 | 0.009 | 0.571 |  |  |  |
| Age >7: Population size (year t-1) | -0.002 | 0.011 | 0.858 |  |  |  |
| D) Number of offspring sired, N=648 observations of 256 males |  |  |  |  |  |  |
| **Intercept** | **-1.694** | **0.371** | **<0.001** | Identity | 0.228 | 0.168-0.308 |
| **Last recorded age** | **0.054** | **0.026** | **0.038** | Measurement | 0.003 | 0.000-0.110 |
| **Age <5** | **0.487** | **0.076** | **<0.001** | year |  |  |
| Age >5 | 0.039 | 0.025 | 0.113 |  |  |  |
| **Population size (year t-1)** | **-0.907** | **0.331** | **0.006** |  |  |  |
| **Age <5: Population size (year t-1)** | **0.159** | **0.071** | **0.027** |  |  |  |
| Age >5: Population size (year t-1) | 0.000 | 0.024 | 0.997 |  |  |  |

### Females: Recruitment success

The best age function model had a threshold at age 6 (CI 4.00-8.03; Table S1D). Recruitment success remained constant before age 6 and declined significantly thereafter (Age_>6_ slope: -0.318±0.043, p<0.001; Table S5B; Figure 2D). Recruitment was higher for female offspring and for mothers with greater last recorded age (offspring sex effect: -0.661±0.090, p<0.001; last recorded age effect: 0.100±0.029, p=0.001; Table S5B). High population size reduced overall recruitment success (- 0.597±0.193, p=0.002; Table S6B) but did not interact with maternal age (Table 4B; Figure 3D). The best fitting model contained a fixed covariate of population size and adding an interaction between age and population size did not significantly improve the fit of the model (Table 2D).

### Females: Offspring birth weight

The best age function model had a threshold at age 7 (CI 6.41-8.45; Table S1E). Offspring birth weight remained constant from maternal ages 4 to 7 and then declined from age 7 onwards (Age_>7_ slope: -0.091±0.010, p<0.001; Table S5C; Figure 2E). Male offspring were generally heavier (0.093 ±0.018, p<0.001), and longer-lived mothers had larger offspring (0.017±0.007, p=0.019; Table S5C). Offspring birth weight decreased with increasing population size (-0.090±0.030, p=0.002; Table S6C), but population size did not interact with declining maternal age (Table 4C; Figure 3E). The best fitting model contained a fixed covariate of population size and adding an interaction between age and population size did not significantly improve the fit of the model (Table 2E).

### Males: Number of offspring sired

The best age function model had a threshold at age 5 (4.00-6.79; Table S1F). Siring success increased significantly between ages 4 and 5, but there was no significant association with age after age 5 (Age_>5_ slope: 0.020±0.025, p=0.423; Table S5D; Figure 2F). Long-lived males sired significantly more offspring (0.079±0.026, p=0.003; Table S5D). Adding the interaction between population size and age significantly improved the fit of this model (Table 2F). This was driven by effects of population size on age related changes in early adulthood, not senescence. Population size interacted with pre-threshold age, with a more rapid increase in performance across ages 4-5 under high density. There was no evidence higher population densities predicted age- related changes in performance in older males (Table 4D; Figure 3F).

## Discussion

Contrary to our predictions, we found no evidence that the rate of senescence in survival or reproductive performance varied with population size in Soay sheep. We further found no support for the hypothesis that demographic senescence is more environmentally sensitive in males than females in a polygynous species. Within our study population, high population size suppressed average adult demographic rates, but did not detectably change age-related declines in either sex. Consistent with previous work, we found no evidence for a sex difference in actuarial senescence and confirmed age-related declines in female breeding probability, offspring recruitment success, and offspring birth weight (Colchero & Clark, 2012; Hayward *et al*., 2015). For males, breeding probability declined with age but offspring number did not. This provides the clearest evidence to date for male reproductive senescence in this system, but it parallels the female pattern: the propensity to breed, rather than offspring number, declines with age. This adds to increasing evidence for asynchrony of ageing patterns among different traits in wild animals, particularly traits related to reproductive success (Hayward *et al*., 2015; Cooper *et al*., 2021; Tsui *et al*., 2025). Why different components of reproduction senesce at different rates remains an important challenge for evolutionary ecologists and our results emphasize that conclusions about reproductive ageing depend strongly on which aspects of reproductive investment and success are measured (Hayward *et al*., 2015; Lemaître & Gaillard, 2017; Moorad & Ravindran, 2022).

Several studies have documented environment-dependent senescence patterns (Reichert *et al*., 2010; Pardo *et al*., 2013) and sex-by-environment-by-age interactions in wild animals (Mysterud *et al*., 2001; Cayuela *et al*., 2021). Human and laboratory studies similarly indicate physiological resilience to stressors declines with age (Kirkland *et al*., 2016; Scheffer *et al*., 2018). There are several possible reasons why we may have failed to detect senescence-environment interactions here. First, we relied on chronological age to test whether senescence renders individuals more demographically vulnerable. However, chronological age is an imperfect indicator of biological state, as individuals vary markedly in their rates of decline in physiological state independent of age. Indeed, trait declines in wild animals are often better predicted by time-to-death than by chronological age (Martin & Festa-Bianchet, 2011; Nussey *et al*., 2011; Hayward *et al*., 2015), mirroring the emergence of biological ageing clocks which predict mortality and morbidity independent of chronological age in humans (Rutledge *et al*., 2022). Such individual variation in biological deterioration, independent of chronological age, could mask effects of the environment on demographic ageing. In future, this could be evaluated using biomarkers of senescence, such as measures of immune function or biological ageing clocks (Peters *et al*., 2019; Le Clercq *et al*., 2023). These biomarkers should predict adult demographic rates more strongly under environmental stress, independent of chronological age. For example, a human epidemiological study found epigenetic ageing rates moderate impacts of environmental risk factors (smoking) on mortality risk (Klopack & Crimmins, 2024). Application of such approaches in wild animals could help understand the drivers of adult demographic variation and improve our ability to predict population and evolutionary dynamics under environmental change.

Apparent late-life declines in demographic rates can reflect both within-individual senescence and between-individual processes like selective disappearance (van de Pol *et al*., 2006; Hayward *et al*., 2013; Nussey *et al*., 2013). Selective disappearance could obscure age-by-environment effects if individuals reaching old age represent a high-quality cohort subset. We detected selective disappearance in reproductive performance models, particularly for males, consistent with other wild animal studies (Bérubé *et al*., 1999; Bouwhuis *et al*., 2009; Hayward *et al*., 2013; Hämäläinen *et al*., 2014). If older individuals are inherently more resilient because they passed through a selective filter earlier in life, this demographic filtering could mask any senescence-mediated losses of resilience. It remains unclear whether inclusion of last recorded age in models fully accounts for these complex effects; simulation studies mapping how within-group centring approaches account for environment-by-age interactions could help resolve this (Van De Pol & Wright, 2009). As individuals age, they gain reproductive and social experience which can counter-balance senescence-associated losses of physiological function. Such experiential ‘buffering’ of late life fitness remains poorly understood in wild systems, but older individuals are increasingly thought to play a central stabilizing role in the ecological dynamics of wild animal systems (McComb *et al*., 2001; Brent *et al*., 2015). As with selective disappearance, this could cause older individuals to appear environmentally resilient due to compensation via behavioural changes linked to experience and knowledge. In both cases, robust biomarkers of physiological function would allow us to separate the within-individual process of deterioration from among-individual processes and within-individual buffering.

Alternatively, weak or absent environmental influences on ageing trajectories may indicate late-life mortality and reproductive failure are primarily driven by intrinsic processes, even in natural populations, with environmental stressors having negligible effects on fundamental functional declines. Current evolutionary theory offers limited insight and prediction as to whether and how age-dependent responses to the environment should evolve and vary under natural conditions. Resource allocation trade-offs underpinning variation in senescence are enshrined in the disposable soma theory of ageing (Kirkwood & Rose, 1991), but exactly how such trade-offs dictate the evolution of life history plasticity and senescence under variable environments remains a topic of considerable debate (Shanley & Kirkwood, 2001; Adler & Bonduriansky, 2014). The ‘developmental theory of ageing’ proposes physiological processes are optimised for early-life growth and survival, rather than later-life function (Maklakov & Chapman, 2019; McAuley, 2024). Such genetically-determined developmental processes may be over- or under-expressed in later life when selection becomes weak, becoming detrimental to physiological function and fitness. Little attention has been paid to how such resource-allocation-independent mechanisms might evolve or respond to variable environments in the evolutionary and ecological literature (Lemaître *et al*., 2020). New evolutionary models addressing how the physiological processes underpinning senescence and phenotypic plasticity should evolve under variable environment are needed to help understand observed differences in environment-dependent senescence among studies.

It is also worth noting we considered only one measure of environmental conditions in our study. Natural environments are complex and multi-dimensional, and different pressures (e.g., food availability, predation, infection risk, weather conditions) may interact with different physiological systems to determine survival and reproductive performance within any study system. These pressures will also vary among populations and species depending on their ecology. This complexity lies at the heart of the challenge of understanding variation in demographic ageing in the wild. Our understanding of the environmental pressures facing Soay sheep on St Kilda is unusually refined: demographic variation is principally influenced by resource competition, winter weather conditions and infection with gastro-intestinal nematodes and we know these pressures have age and sex dependent effects (Gulland, 1992; Catchpole *et al*., 2000; Coulson *et al*., 2001). Here, we tested interactions between an indicator of just one of these pressures (resource competition) and ageing. Considering a wider range of environmental metrics and biomarkers of specific aspects of physiological function, such as measures of immune function or combined measures of environmental adversity (e.g., Tung *et al*., 2016; Ortiz-Ross & Blumstein, 2024), might reveal changes in age-specific resilience a single environmental measure cannot.

Increasing demographic environmental sensitivity, due to loss of physiological resilience, in later life could impact how wild populations respond to environmental change and underpin the evolution of variation in senescence and lifespan. Our study provides an unusually detailed test of this hypothesis in a wild mammal and shows demographic senescence is not dependent on prevailing levels of resource competition in either sex. Our findings challenge the prediction that older animals should be more sensitive to environmental conditions, but also highlight the need for biomarkers of underlying physiological function, measures of experiential gain and compensation and methods to account for selective disappearance under variable environments. Our work also demonstrates the current gap in our theoretical knowledge surrounding the evolution of age-dependent phenotypic plasticity and physiological resilience. Our results illustrate an emerging picture in which a simplistic ‘males are always more vulnerable’ rule does not apply universally. In some systems, males may indeed show heightened sensitivity to environmental stress, whereas in others sex differences may be small or absent, or appear only under specific stressors. Similar studies across wild populations should consider that sex-specific ageing and environmental sensitivity may depend on the interactions between sex-specific physiology and reproductive schedules, the social structure of populations, and which ecological challenges dominate variation in late-life survival and reproduction in specific populations.

## Data and code availability statement

The data and code associated with this data is available at [Dryad link here]. Supplementary material is available online.

## Author contributions

E.D.D.: conceptualization, data curation, formal analysis, methodology, visualization, writing-original draft, writing-review and editing; X.B.: investigation, data curation, writing—review and editing; J.M.P.: funding acquisition, investigation, project administration; J.G.P.: investigation, data curation, writing—review and editing; D.H.N.: conceptualization, funding acquisition, project administration, supervision, writing—original draft, writing—review and editing; H.F.: conceptualization, data curation, formal analysis, methodology, supervision, writing—original draft, writing— review and editing. All authors gave final approval for publication and agreed to be held accountable for the work performed therein.

## Acknowledgements

We are grateful to everyone who has been involved in the long-term study of Soay sheep on St Kilda, Scotland, over the last 39 years, particularly Ian Stevenson, Michael Morrisey, Jon Slate, Susan Johnston, Tim Clutton-Brock, Loeske Kruuk and many other researchers and fieldworkers. We are also grateful to the National Trust for Scotland for permissions to work on St Kilda, and QinetiQ and Kilda Cruises for logistical assistance in the field.

