## Supplementary Material for "No evidence sex-specific senescence varies with resource competition in Soay sheep"

### Tables

*Table S1. Summary of the thresholds and AICs of the threshold models tested for: (A) binomial GLMMs of survival, B) binomial GLMMs of breeding probability, (C) binomial GLMMs of twinning probability, (D) binomial GLMMs of offspring recruitment, (E) gaussian GLMMs of offspring birth weight, and (F) negative binomial GLMMs of number of offspring sired. Models either had no/a single threshold. The final models chosen, the best age function models, are in bold.*

| Threshold tested | AIC | ∆AIC |
|---|---|---|
| A) Survival | | |
| **No threshold** | **3517.784** | **0.571** |
| 4 | 3520.950 | 3.737 |
| 5 | 3521.776 | 4.563 |
| 6 | 3521.269 | 4.056 |
| 7 | 3519.700 | 2.487 |
| 8 | 3517.213 | 0.000 |
| B) Breeding probability | | |
| No threshold | 3549.570 | 30.954 |
| 5 | 3533.179 | 14.562 |
| 6 | 3522.408 | 3.791 |
| 7 | 3518.933 | 0.316 |
| **8** | **3518.617** | **0.000** |
| 9 | 3531.952 | 13.335 |
| C) Twinning probability | | |
| No threshold | 2581.150 | 11.061 |
| 5 | 2576.118 | 6.028 |
| **6** | **2570.089** | **0.000** |
| 7 | 2574.203 | 4.114 |
| 8 | 2578.482 | 8.393 |
| 9 | 2579.361 | 9.271 |
| D) Recruitment probability | | |
| No threshold | 3846.808 | 5.153 |
| 5 | 3842.074 | 0.419 |
| **6** | **3841.655** | **0.000** |
| 7 | 3842.310 | 0.656 |
| 8 | 3843.639 | 1.984 |
| 9 | 3845.601 | 3.947 |

| | | |
|---|---|---|
| E) Offspring birth weight | | |
| No threshold | 4232.436 | 20.575 |
| 5 | 4220.200 | 8.340 |
| 6 | 4217.568 | 5.707 |
| **7** | **4211.861** | **0.000** |
| 8 | 4212.161 | 0.301 |
| 9 | 4219.149 | 7.289 |
| F) Number of offspring sired | | |
| No threshold | 2825.875 | 17.189 |
| **5** | **2808.686** | **0.000** |
| 6 | 2810.244 | 1.558 |
| 7 | 2812.258 | 3.572 |
| 8 | 2818.681 | 9.995 |
| 9 | 2817.595 | 8.910 |

*Table S2. Summary of the thresholds and AICs of the threshold models tested for GLMMs of breeding probability for either (A) only females, or (B) only males. Models either had no/a single threshold. The lowest AIC models are in bold.*

| **Threshold tested** | **AIC** | **∆AIC** |
|---|---|---|
| A) Females | | |
| No threshold | 2136.629 | 19.257 |
| 5 | 2125.269 | 7.897 |
| 6 | 2119.438 | 2.066 |
| **7** | **2117.372** | **0.000** |
| 8 | 2119.111 | 1.739 |
| 9 | 2122.403 | 5.031 |
| B) Males | | |
| No threshold | 1407.067 | 10.134 |
| 5 | 1401.862 | 4.929 |
| 6 | 1398.522 | 1.588 |
| 7 | 1397.646 | 0.713 |
| **8** | **1396.933** | **0.000** |
| 9 | 1404.290 | 7.357 |

*Table S3. Summary and fixed and random effects of binomial GLMMs of survival; (A) best age function model, (B) best age function model including a fixed effect of population size in year t, and (C) best age function model including 2-way interactions between age, sex and population size in year t. Significant effects in bold. $N_{female}$=3604 obvs of 649 individuals; $N_{male}$=1097 obvs of 371 individuals.*

| **Fixed effects** | **Estimate** | **Std. error** | **p-value** | **Random effects** | **Variance** | **95% CI** |
|---|---|---|---|---|---|---|
| A) Best age function model | | | | | | |
| **Intercept** | **4.705** | **0.374** | **<0.001** | Identity | 0.096 | 0.005-1.695 |
| **Sex (male)** | **-1.879** | **0.300** | **<0.001** | Birth year | 0.019 | 0.002-0.158 |
| **Age** | **-0.440** | **0.042** | **<0.001** | Measurement | 2.284 | 1.332-3.917 |
| Age:Sex (male) | 0.038 | 0.057 | 0.505 | year | | |
| B) Best age function model including a fixed effect of population size in year t | | | | | | |
| **Intercept** | **4.496** | **0.358** | **<0.001** | Identity | 0.079 | 0.003-2.341 |
| **Sex (female)** | **-1.872** | **0.299** | **<0.001** | Birth year | 0.019 | 0.002-0.153 |
| **Age** | **-0.433** | **0.041** | **<0.001** | Measurement | 1.797 | 1.020-3.166 |
| **Population size in year t** | **-0.641** | **0.233** | **0.006** | year | | |
| Age:Sex (male) | 0.040 | 0.057 | 0.478 | | | |
| C) Best age function model including 2-way interactions between age, sex and population size in year t | | | | | | |
| **Intercept** | **4.449** | **0.361** | **<0.001** | Identity | 0.069 | 0.001-3.346 |
| **Sex (female)** | **-1.864** | **0.302** | **<0.001** | Birth year | 0.021 | 0.003-0.148 |
| **Age** | **-0.425** | **0.042** | **<0.001** | Measurement | 1.747 | 0.988-3.090 |
| Population size in year t | -0.526 | 0.279 | 0.059 | year | | |
| Age:Sex (male) | 0.042 | 0.057 | 0.456 | | | |
| Age:Population size in year t | -0.019 | 0.026 | 0.452 | | | |
| Sex (male): Population size in year t | -0.036 | 0.121 | 0.769 | | | |

*Table S4. Summary and fixed and random effects of binomial GLMMs of breeding probability; (A) best age function model after model selection, (B) best age function model including a fixed effect of population size in year t, and (C) best age function model including 2-way interactions between age, sex and population size in year t. Significant effects in bold. $N_{female}$=3774 obvs of 807 individuals; $N_{male}$=1287 obvs of 418 individuals.*

| **Fixed effects** | **Estimate** | **Std. error** | **p-value** | **Random effects** | **Variance** | **95% CI** |
|---|---|---|---|---|---|---|
| A) Best age function model | | | | | | |
| **Intercept** | **4.414** | **0.581** | **<0.001** | Identity | 3.449 | 2.674-4.447 |
| **Sex (male)** | **-7.644** | **0.777** | **<0.001** | Measurement year | 2.975 | 1.715-5.159 |
| Last recorded age | -0.081 | 0.050 | 0.108 | | | |
| Age <8 | -0.056 | 0.057 | 0.327 | | | |
| **Age >8** | **-0.525** | **0.077** | **<0.001** | | | |
| **Sex (male): Last recorded age** | **0.446** | **0.096** | **<0.001** | | | |
| **Age <8: Sex (male)** | **0.195** | **0.094** | **0.038** | | | |
| Age >8: Sex (male) | -0.497 | 0.287 | 0.084 | | | |
| B) Best age function model including a fixed effect of population size in year t-1 | | | | | | |
| **Intercept** | **4.355** | **0.582** | **<0.001** | Identity | 3.447 | 2.673-4.445 |
| **Sex (male)** | **-7.644** | **0.777** | **<0.001** | Measurement year | 2.908 | 1.675-5.049 |
| Last recorded age | -0.082 | 0.050 | 0.103 | | | |
| Age <8 | -0.053 | 0.057 | 0.350 | | | |
| **Age >8** | **-0.524** | **0.077** | **<0.001** | | | |
| Population size | -0.224 | 0.263 | 0.394 | | | |
| **Sex (male): Last recorded age** | **0.446** | **0.096** | **<0.001** | | | |
| **Age <8: Sex (male)** | **0.195** | **0.094** | **0.038** | | | |
| Age >8: Sex (male) | -0.498 | 0.287 | 0.083 | | | |
| C) Best age function model including 2-way interactions between age, sex and population size in year t-1 | | | | | | |
| **Intercept** | **4.345** | **0.582** | **<0.001** | Identity | 3.444 | 2.671-4.442 |
| **Sex (female)** | **-7.602** | **0.781** | **<0.001** | Measurement year | 2.911 | 1.676-5.055 |
| Last recorded age | -0.082 | 0.050 | 0.102 | | | |
| Age <8 | -0.052 | 0.057 | 0.366 | | | |
| **Age >8** | **-0.528** | **0.077** | **<0.001** | | | |
| Population size in year t-1 | -0.339 | 0.359 | 0.346 | | | |
| **Sex (male): Last recorded age** | **0.445** | **0.097** | **<0.001** | | | |
| Age <8: Sex (male) | 0.188 | 0.097 | 0.052 | | | |
| Age >8: Sex (male) | -0.492 | 0.287 | 0.087 | | | |
| Age <8: Population size in year t-1 | 0.022 | 0.044 | 0.616 | | | |
| Age >8: Population size in year t-1 | -0.039 | 0.072 | 0.589 | | | |
| Sex (male): Population size in year t-1 | 0.013 | 0.127 | 0.921 | | | |

*Table S5. Summary and fixed and random effects of GLMMs best age function models for (A) twinning probability (binomial; threshold at age 6), (B) recruitment success (binomial; threshold at age 6; reference level female offspring), (C) offspring birth weight (gaussian; threshold at age 7; reference level female offspring) and (D) number of offspring sired by a male (negative binomial; threshold age at 5). Significant effects in bold.*

| **Fixed effects** | **Estimate** | **Std. error** | **p-value** | **Random effects** | **Variance** | **95% CI** |
|---|---|---|---|---|---|---|
| A) Female twinning probability, N=3155 obvs of 737 females | | | | | | |
| **Intercept** | **-4.584** | **0.563** | **<0.001** | Identity | 2.236 | 1.651-3.029 |
| Last recorded age | -0.012 | 0.044 | 0.793 | Measurement | 0.107 | 0.041-0.277 |
| **Age <6** | **0.435** | **0.089** | **<0.001** | year | | |
| Age >6 | 0.016 | 0.050 | 0.753 | | | |
| B) Recruitment success N=3638 obvs of 738 females | | | | | | |
| Intercept | 0.212 | 0.473 | 0.654 | Identity | 0.626 | 0.436-0.899 |
| **Last recorded age** | **0.100** | **0.029** | **0.001** | Measurement | 1.927 | 1.162-3.196 |
| **Offspring sex (male)** | **-0.661** | **0.090** | **<0.001** | year | | |
| Age <6 | -0.080 | 0.064 | 0.212 | | | |
| **Age >6** | **-0.318** | **0.043** | **<0.001** | | | |
| C) Offspring birth weight, N=2881 obvs of 707 females | | | | | | |
| **Intercept** | **1.791** | **0.083** | **<0.001** | Identity | 0.092 | 0.078-0.109 |
| **Last recorded age** | **0.017** | **0.007** | **0.019** | Measurement | 0.041 | 0.024-0.069 |
| **Offspring capture age (days)** | **0.115** | **0.004** | **<0.001** | year | | |
| **Offspring sex (male)** | **0.093** | **0.018** | **<0.001** | | | |
| Age <7 | -0.015 | 0.009 | 0.094 | | | |
| **Age >7** | **-0.091** | **0.010** | **<0.001** | | | |
| D) Offspring sired model, N=690 obvs of 277 males | | | | | | |
| **Intercept** | **-1.363** | **0.358** | **<0.001** | Identity | 0.228 | 0.168-0.309 |
| **Last recorded age** | **0.079** | **0.026** | **0.003** | Measurement | 0.022 | 0.007-0.070 |
| **Age <5** | **0.388** | **0.072** | **<0.001** | year | | |
| Age >5 | 0.020 | 0.025 | 0.423 | | | |

*Table S6. Summary and fixed and random effects of GLMMs best age function models, including a fixed effect of population size, for (A) twinning probability (binomial; threshold at age 6), (B) recruitment success (binomial; threshold at age 6; reference level female offspring), (C) offspring birth weight (gaussian; threshold at age 7; reference level female offspring) and (D) number of offspring sired by a male (negative binomial; threshold age at 5). Significant effects in bold.*

| **Fixed effects** | **Estimate** | **Std. error** | **p-value** | **Random effects** | **Variance** | **95% CI** |
|---|---|---|---|---|---|---|
| A) Female twinning probability, N=3155 obvs of 737 females | | | | | | |
| **Intercept** | **-4.622** | **0.561** | **<0.001** | Identity | 2.244 | 1.658-3.037 |
| Last recorded age | -0.024 | 0.044 | 0.591 | Measurement | 0.064 | 0.021-0.192 |
| **Age <6** | **0.455** | **0.089** | **<0.001** | year | | |
| Age >6 | 0.023 | 0.050 | 0.639 | | | |
| **Population size t-1** | **-0.339** | **0.077** | **<0.001** | | | |
| B) Recruitment success N=3638 obvs of 738 females | | | | | | |
| Intercept | 0.063 | 0.461 | 0.891 | Identity | 0.624 | 0.435-0.896 |
| **Last recorded age** | **0.095** | **0.029** | **0.001** | Measurement | 1.468 | 0.876-2.461 |
| **Offspring sex (male)** | **-0.661** | **0.090** | **<0.001** | year | | |
| Age <6 | -0.072 | 0.064 | 0.259 | | | |
| **Age >6** | **-0.313** | **0.043** | **<0.001** | | | |
| **Population size t** | **-0.597** | **0.193** | **0.002** | | | |
| C) Offspring birth weight, N=2881 obvs of 707 females | | | | | | |
| **Intercept** | **1.779** | **0.081** | **<0.001** | Identity | 0.092 | 0.078-0.109 |
| **Last recorded age** | **0.017** | **0.007** | **0.025** | Measurement | 0.031 | 0.018-0.053 |
| **Offspring capture age (days)** | **0.115** | **0.004** | **<0.001** | year | | |
| **Offspring sex (male)** | **0.093** | **0.018** | **<0.001** | | | |
| Age <7 | -0.013 | 0.009 | 0.139 | | | |
| **Age >7** | **-0.090** | **0.010** | **<0.001** | | | |
| **Population size t-1** | **-0.090** | **0.030** | **0.002** | | | |
| D) Offspring sired model, N=690 obvs of 277 males | | | | | | |
| **Intercept** | **-1.398** | **0.348** | **<0.001** | Identity | 0.231 | 0.171-0.312 |
| **Last recorded age** | **0.060** | **0.026** | **0.021** | Measurement | 0.003 | 0.000-0.239 |
| **Age <5** | **0.416** | **0.070** | **<0.001** | year | | |
| Age >5 | 0.045 | 0.025 | 0.066 | | | |
| **Population size t-1** | **-0.154** | **0.031** | **<0.001** | | | |

### Figures

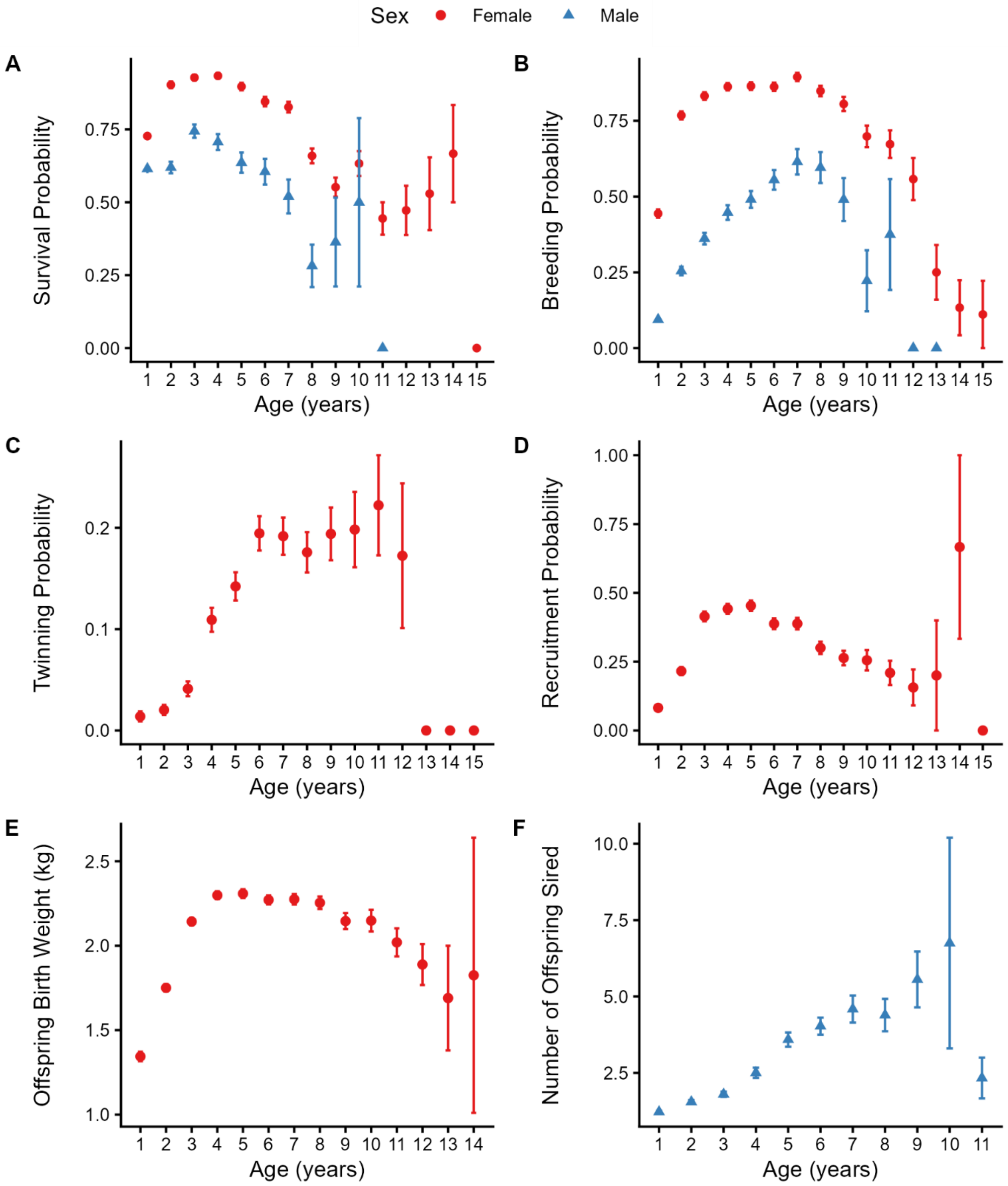

*Figure S1. Plots of raw data across the adult age range for male and female (A) survival probability and (B) breeding probability; female (C) twinning probability, (D) recruitment probability, (E) offspring birth wight; and (F) the number of offspring sired by males in Soay sheep. Red circular and blue triangular points and error bars represent the mean and associated standard error at each age for females and males, respectively.*
